# Discovery of flavivirus-derived endogenous viral elements in two *Anopheles* mosquito genomes supports the existence of *Anopheles*-associated insect-specific flaviviruses

**DOI:** 10.1101/064907

**Authors:** Sebastian Lequime, Louis Lambrechts

## Abstract

The *Flavivirus* genus encompasses several arboviruses of public health significance such as dengue, yellow fever, and Zika viruses. It also includes insect-specific flaviviruses (ISFs) that are only capable of infecting insect hosts. The vast majority of mosquito-infecting flaviviruses have been associated with mosquito species of the *Aedes* and *Culex* genera in the Culicinae subfamily, which also includes most arbovirus vectors. Mosquitoes of the Anophelinae subfamily are not considered significant arbovirus vectors, however flaviviruses have occasionally been detected in field-caught *Anopheles* specimens. Whether such observations reflect occasional spillover or laboratory contamination or whether *Anopheles* mosquitoes are natural hosts of flaviviruses is unknown. Here, we provide *in silico* and *in vivo* evidence of transcriptionally active, flavivirus-derived endogenous viral elements (EVEs) in the genome of *Anopheles minimus* and *Anopheles sinensis*. Such non-retroviral endogenization of RNA viruses is consistent with a shared evolutionary history between flaviviruses and *Anopheles* mosquitoes. Phylogenetic analyses of the two newly described EVEs support the existence of a distinct clade of *Anopheles*-associated ISFs.

## Introduction

Flaviviruses are positive-sense, single-stranded RNA viruses that infect a broad range of hosts including vertebrates (e.g., birds, primates) and arthropods (e.g., ticks, mosquitoes). In addition to major arboviruses of public health significance such as dengue, Zika, West Nile and yellow fever viruses, the *Flavivirus* genus also includes vertebrate-specific (not known vector; NKV) and insect-specific (insect-specific flaviviruses; ISFs) members (Moureau et al. 2015). The majority of mosquito-infecting flaviviruses have been associated with members of the Culicinae subfamily, mainly from the *Culex* and *Aedes* genera. *Anopheles* mosquitoes in the Anophelinae subfamily are well known for their role in the transmission of human malaria parasites, but they are not considered important vectors of arboviruses in general, and of flaviviruses in particular. Nevertheless, several studies have detected flaviviruses in field-caught *Anopheles* mosquitoes from different parts of the world. In North America, West Nile virus (WNV) was detected in *Anopheles punctipennis* (Bernard et al. 2001; Kulasekera et al. 2001). In Asia, Japanese encephalitis virus was detected in *Anopheles sinensis* (Feng et al. 2012; Liu et al. 2013). In Europe, *Anopheles maculipennis* was found positive for WNV (Filipe 1972; Kemenesi et al. 2014), Usutu virus (Calzolari et al. 2013) and Batai virus (Calzolari et al. 2010). Interestingly, some ISFs were also detected in *An. sinensis* (Zuo et al. 2014; Liang et al. 2015) and *Anopheles atroparvus* (Aranda et al. 2009). It is unknown whether these detections reflect occasional spillover or laboratory contamination, or whether *Anopheles* mosquitoes are in fact natural hosts of flaviviruses.

Endogenous viral elements (EVEs) are chromosomal integrations of partial or full viral genetic material into the genome of a host species. Not only retroviruses, whose replication cycle includes incorporation of a DNA form of the RNA viral genome into the host cell genome, but virtually all types of eukaryotic viruses can become endogenous (Feschotte and Gilbert 2012). Non-retroviral EVEs have been documented in the genome of a wide variety of host species, including vertebrates and arthropods (Feschotte and Gilbert 2012). Unlike detection of exogenous viruses, subject to possible laboratory or environmental contamination, EVEs are likely to reflect a long-lasting evolutionary relationship between an RNA virus and its natural host. This is because endogenization, for a single-stranded RNA virus, requires (1) reverse transcription, (2) integration of virus-derived DNA into the genome of germinal host cells and (3) fixation of the integrated sequence in the host population (Holmes 2011; Aiewsakun and Katzourakis 2015). The low probability of this combination of events makes endogenization exceedingly unlikely to occur unless the viral infection is common in the host population over long evolutionary times. For example, flavivirus-derived EVEs have been reported in the genome of *Aedes aegypti* (Crochu et al. 2004; Katzourakis and Gifford 2010) and *Aedes albopictus* (Roiz et al. 2009; Chen et al. 2015). These EVEs are phylogenetically related to the clade of *Aedes*-associated ISFs (Crochu et al. 2004; Roiz et al. 2009; Katzourakis and Gifford 2010; Chen et al. 2015), which is consistent with the ancient evolutionary relationship between *Aedes* mosquitoes and ISFs.

Here, we report the discovery of two flavivirus-derived EVEs in the genomes of *An*. *minimus* and *An. sinensis* mosquitoes. We screened 24 publicly available *Anopheles* genomes (Holt et al. 2002; Zhou et al. 2014; Logue et al. 2015; Neafsey et al. 2015) for flaviviral sequences, and validated *in silico* hits both at the DNA and RNA levels *in vivo*. The two newly described flavivirus-derived EVEs are phylogenetically related to ISFs, and support the existence of a previously unsuspected *Anopheles*-associated clade of ISFs.

## Material and Methods

### 1. *In silico* survey

#### 1.1. Genome screen

Twenty-four *Anopheles* genomes (full list and accession numbers are provided in Table 1) were screened by tBLASTn (BLAST+ 2.2.28) (Camacho et al. 2009) for the presence of flavivirus-derived EVEs using a collection of 50 full flavivirus polyprotein queries (full list and accession numbers are provided in Table S1) representing the currently known diversity of the *Flavivirus* genus. The sequences of hits whose E-value was > E^−4^ were extracted using an in-house bash shell script. In order to reconstruct putative flavivirus-derived EVEs, BLAST hits were clustered using CD-HIT v.4.6.1 (Li and Godzik 2006) and aligned using MAFFT v7.017 (Katoh et al. 2002). Putative EVEs were extracted and used as query for a reciprocal tBLASTn (BLAST+ 2.2.30) against the National Center for Biotechnology Information (NCBI) nucleotide database (E-value > 10^−4^). Genetic features of identified EVE were analyzed using the NCBI Conserved Domain Database (Marchler-Bauer et al. 2015). Nucleotide sequence data reported are available in the Third Party Annotation Section of the DDBJ/ENA/GenBank databases under the accession numbers TPA: BK009978-BK009980.

**Table 1:**
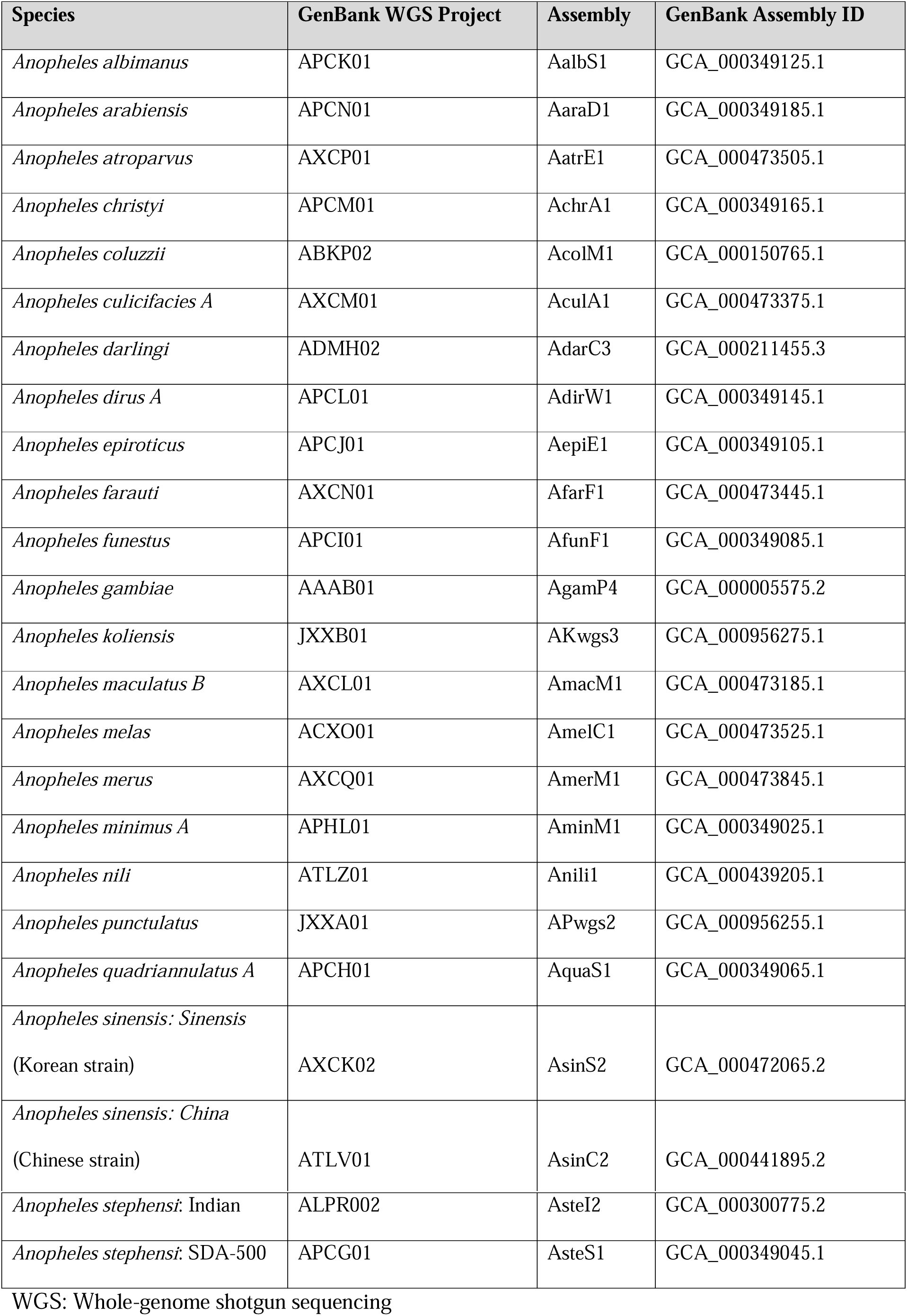
*Anopheles* genomes screened in this study.

**Table 2:** Newly described flavivirus-derived EVEs in *An. minimus* and *An. sinensis* genomes.

| Element | GenBank<br>accession no. | Protein<br>identity | Supercontig<br>accession no. | Super<br>contig<br>length<br>(bp) | Coordinates |  |  |  |
| --- | --- | --- | --- | --- | --- | --- | --- | --- |
|  |  |  |  |  | Supercontig |  | Viral genome* |  |
|  |  |  |  |  | Start | End | Start | End |
| *An. minimus* EVE | BK009978 | NS4-NS5 | KB664005.1 | 2,043 | 107 | 1,987 | 6,567 | 8,432 |
| *An. sinensis* EVE<br>(Korean strain) | BK009979 | NS3 | AXCK0202474<br>4 | 2,797 | 822 | 1,613 | 5,214 | 5,822 |
| *An. sinensis* EVE<br>(Chinese strain) | BK009980 | NS3 | ATLV01019207<br>.1 | 7,380 | 3,578 | 4,376 | 5,214 | 5,822 |

\* Viral genomes coordinates are based on the closest tBLASTn hits: Nienokoue virus
(GenBank accession no. JQ957875) for the *An. minimus* EVE and *Culex* flavivirus (GenBank
accession no. JQ308188) for the *An. sinensis* EVE.

#### 1.2. Phylogenetic analyses

Translated EVE sequences were aligned to the corresponding sections of several flavivirus polyproteins (Table S1) with MAFFT v7.017 and phylogenetically uninformative positions were trimmed using TrimAI v.1.3 (Capella-Gutiérrez et al. 2009) accessed through the webserver Phylemon 2 (Sánchez et al. 2011). The trimmed alignments were used to construct phylogenetic trees with PhyML Best AIC Tree (Sánchez et al. 2011). Best substitution models were Blosum62+I+G+F for *An. minimus* and Blosum62+I+G for *An*. *sinensis*.

#### 1.3. Transcriptome screen

Published RNA sequencing (RNA-seq) data were retrieved from NCBI Sequence Read Archive (Leinonen et al. 2011) and explored for the presence of previously identified EVE sequences. Only one *An. minimus* transcriptome sequence read archive was found under accession number SRX265162. Six *An. sinensis* transcriptome sequence read archives were found under accession numbers SRX448985, SRX449003, SRX449006 and SRX277584 for experiments using Illumina sequencing technology, and SRX218691 and SRX218721 for experiments using Roche 454 sequencing technology. RNA-seq reads were mapped to the EVE nucleotide sequence using Bowtie2 v2.1.0 (Langmead and Salzberg 2012). The alignment file was converted, sorted and indexed with Samtools v0.1.19 (Li et al. 2009). Coverage was assessed using bedtools v2.17.0 (Quinlan and Hall 2010).

### 2.2 *In vivo* validation

#### 2.1. Mosquitoes

*Anopheles minimus* and *An. sinensis* mosquitoes were obtained through BEI Resources (www.beiresources.org), National Institute of Allergy and Infectious Diseases, National Institutes of Health (*An. minimus* MINIMUS1, MRA-729; *An. sinensis* SINENSIS, MRA-1154). *Anopheles minimus* and *An. sinensis* specimens came from the 132^nd^ and 65^th^ generations of laboratory colonization, respectively. Eggs were hatched in filtered tap water, reared in 24×34×9 cm plastic trays and fed with fish food (TetraMin, Tetra, Melle, Germany). Adults were maintained in 30×30×30 cm screened cages under controlled insectary conditions (28±1°C, 75±5% relative humidity, 12:12 hour light-dark cycle). They were provided with cotton soaked in a 10% (m/v) sucrose solution *ad libitum*. *Anopheles stephensi* nucleic acids, used as a reaction control, were kindly provided by the Genetics and Genomics of Insect Vectors unit, Institut Pasteur, Paris.

#### 2.2. EVE genomic integration

Mosquitoes were homogenized in pools of 10 separated by sex in 300 µL of Dulbecco’s phosphate-buffered saline (DPBS) during two rounds of 30 sec at 5,000 rpm in a mixer mill (Precellys 24, Bertin Technologies, Montigny le Bretonneux, France). DNA was extracted using All Prep DNA/RNA Mini Kit (Qiagen, Hilden, Germany) following the manufacturer’s instructions. EVE presence in genomic DNA was assessed by 35 cycles of PCR using Taq Polymerase (Invitrogen, Thermo Fisher Scientific, Waltham, MA, USA) (Table S2). PCR primers were designed to generate an amplicon spanning part of the EVE sequence and a section of the flanking host sequence. Identity of the EVE sequence was confirmed by Sanger sequencing of the PCR product.

#### 2.3. EVE transcription level

Mosquitoes were homogenized in pools of 5 separated by sex or development stage in 300 µL of DPBS during two rounds of 30 sec at 5,000 rpm in a mixer mill (Precellys 24). RNA was extracted from mosquito homogenates separated by sex using TRIzol Reagent (Life Technologies, Thermo Fisher Scientific, Waltham, MA, USA) following the manufacturer’s instructions. Samples were treated with Turbo DNA-free kit (Life Technologies) and reverse transcribed using random hexamers and M-MLV reverse transcriptase (Invitrogen). Complementary DNA was amplified with 35 cycles of PCR for *An. minimus* and 40 cycles of PCR for *An. sinensis,* respectively, using DreamTaq polymerase (Thermo Fisher Scientific) and primers located within the EVE sequence (Table S2). To verify that RNA samples were free of DNA contamination, two sets of primers spanning exons 3 and 4 of the RPS7 gene of both *Anopheles* species (under VectorBase annotation number AMIN008193 and ASIC017918 for *An. minimus* and *An. sinensis*, respectively) were designed (Table S2). Because the corresponding DNA sequence includes intron 3, DNA contamination is expected to result in a larger PCR product. The length of intron 3 is 252 base pairs (bp) for *An. minimus* and 295 bp for *An. sinensis*.

## Results

The *in silico* screen of 24 *Anopheles* genomes identified two flavivirus-derived EVEs, one in the reference genome sequence of *An. minimus* and one in both genome sequences available for *An. sinensis* (Table 1).

### *An. minimus* EVE

The *An. minimus* EVE is 1,881 bp long (627 amino acid residues) with Nienokoue virus as the closest BLAST hit (44% amino-acid identity). The integrated sequence spans non-structural protein 4A (NS4A), NS4B and NS5 (Figure 1A) without any stop codon. Conserved domain search identified the NS5-methyltransferase domain involved in RNA capping and part of the RNA-directed RNA-polymerase domain. Phylogenetically, the *An*. *minimus* EVE is sister to the ISF clade (Figure 1B). The *An. minimus* genome supercontig where the EVE was detected is 2,043 bp long and consists almost exclusively of the EVE sequence.

**Figure 1:**
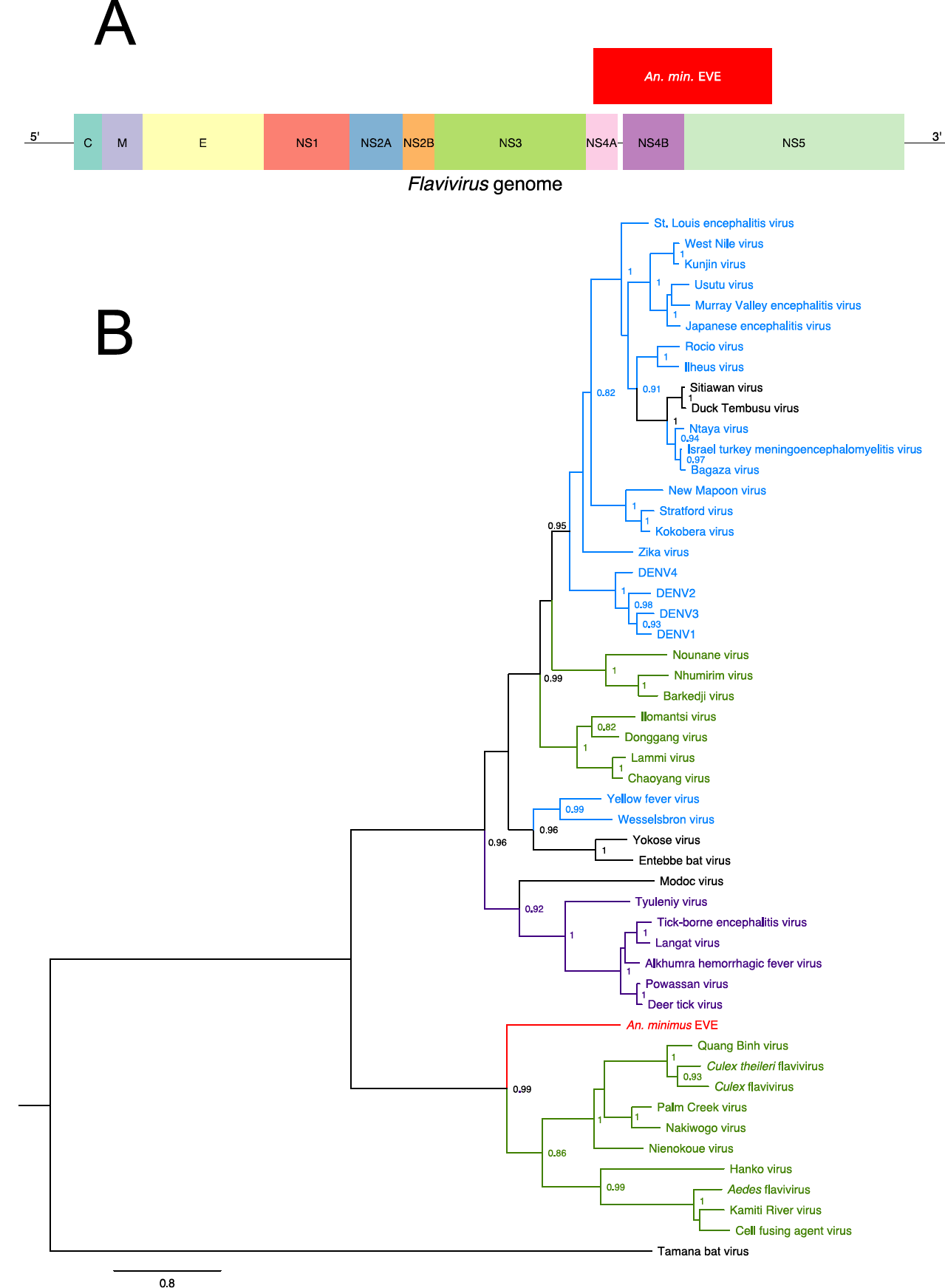
Discovery of a flavivirus-derived EVE in *An. minimus*. (A) EVE location in a generic *Flavivirus* genome. Positioning is based on the genome sequence of Nienokoue virus (GenBank accession no. JQ957875). C=capsid protein, E=envelope glycoprotein, M=membrane glycoprotein, NS1=non-structural glycoprotein 1; NS2A=non-structural protein 2A; NS2B= non-structural protein 2B; NS3=non-structural protein 3 (protease/helicase); NS4A=non-structural protein 4A; NS4B=non-structural protein 4B; NS5=non-structural protein 5 (RNA-dependent RNA polymerase). (B) Phylogenetic relationships of the newly discovered *An. minimus* flavivirus-derived EVE with exogenous flaviviruses. Maximum likelihood trees were constructed based on the translated EVE sequence. Clades are color-coded according to known host specificity: green, ISFs; purple, tick-borne arboviruses; black, ‘not known vector’ (vertebrate specific); blue, mosquito-borne arboviruses; red: EVEs. Scale bar indicates the number of substitutions. Node values represent Shimodaira-Hasegawa (SH)-like branch support (only values > 0.8 are shown).

Presence of the EVE was verified *in vivo* by PCR on genomic DNA (Figure 2A), followed by amplicon sequencing to confirm identity. The *An. minimus* EVE was found in both male and female genomic DNA, and was transcriptionally expressed for all combinations of sex and development stages tested (Figure 2B). Evidence for transcriptional activity of the *An*. *minimus* EVE was confirmed in published RNA-seq data (Figure S1A).

**Figure 2:**
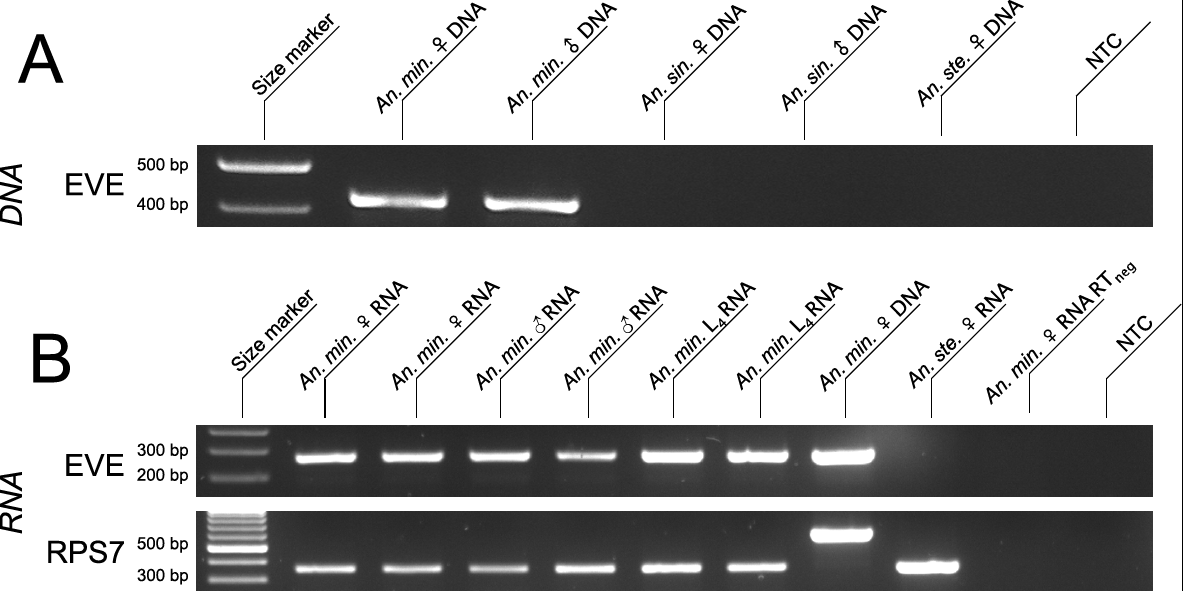
*In vivo* detection of the *An. minimus* flavivirus-derived EVE. (A) EVE detection in genomic DNA. Lane 1: size marker; lane 2: amplified genomic DNA from a pool of 10 *An*. *minimus* adult females; lane 3: amplified genomic DNA from a pool of 10 *An. minimus* adult males; lane 4: amplified genomic DNA from a pool of 10 *An. sinensis* adult females; lane 5: amplified genomic DNA from a pool of 10 *An. sinensis* adult males; lane 6: amplified DNA from a pool of 10 *An. stephensi* females; lane 7: no template control (NTC). (B) EVE detection in total RNA. Lane 1: size marker; lanes 2 and 3: amplified cDNA from pools of 5 adult females; lanes 4 and 5: amplified cDNA from pools of 5 adult males; lanes 6 and 7: amplified cDNA from pools of 5 L_4_ larvae; lane 8: amplified DNA from a pool of 10 females; lane 9: amplified cDNA from a pool of 5 *An. stephensi* females; lane 10: DNA contamination control (no reverse transcription) using the same pool of 5 adult females as lane 2; lane 11: no template control (NTC). First row: EVE; second row: RPS7 (control gene). The RPS7 target DNA sequence includes an intron, so that DNA contamination is expected to result in a larger PCR product.

### *An. sinensis* EVE

The *An. sinensis* EVE was detected in two distinct genome sequences that are available for this mosquito species, derived from a Korean and a Chinese strain, respectively. The EVE is 792 bp long (264 amino acid residues) and 799 bp long (266 amino acid residues) for the Korean and Chinese strains, respectively (Table 1). The closest BLAST hit is *Culex* flavivirus (45% amino-acid identity for the Korean strain, 44% amino-acid identity for the Chinese strain). The integrated sequence corresponds to the middle part of NS3 (Figure 3A) and contains six and eight stop codons in the Korean and Chinese strains, respectively. Conserved domain search identified the presence of a P-loop containing the nucleoside triphosphate hydrolase domain found in the NS3 protein of exogenous flaviviruses. Phylogenetically, the *An. sinensis* EVE is sister to the ISF clade (Figure 3B).

**Figure 3:**
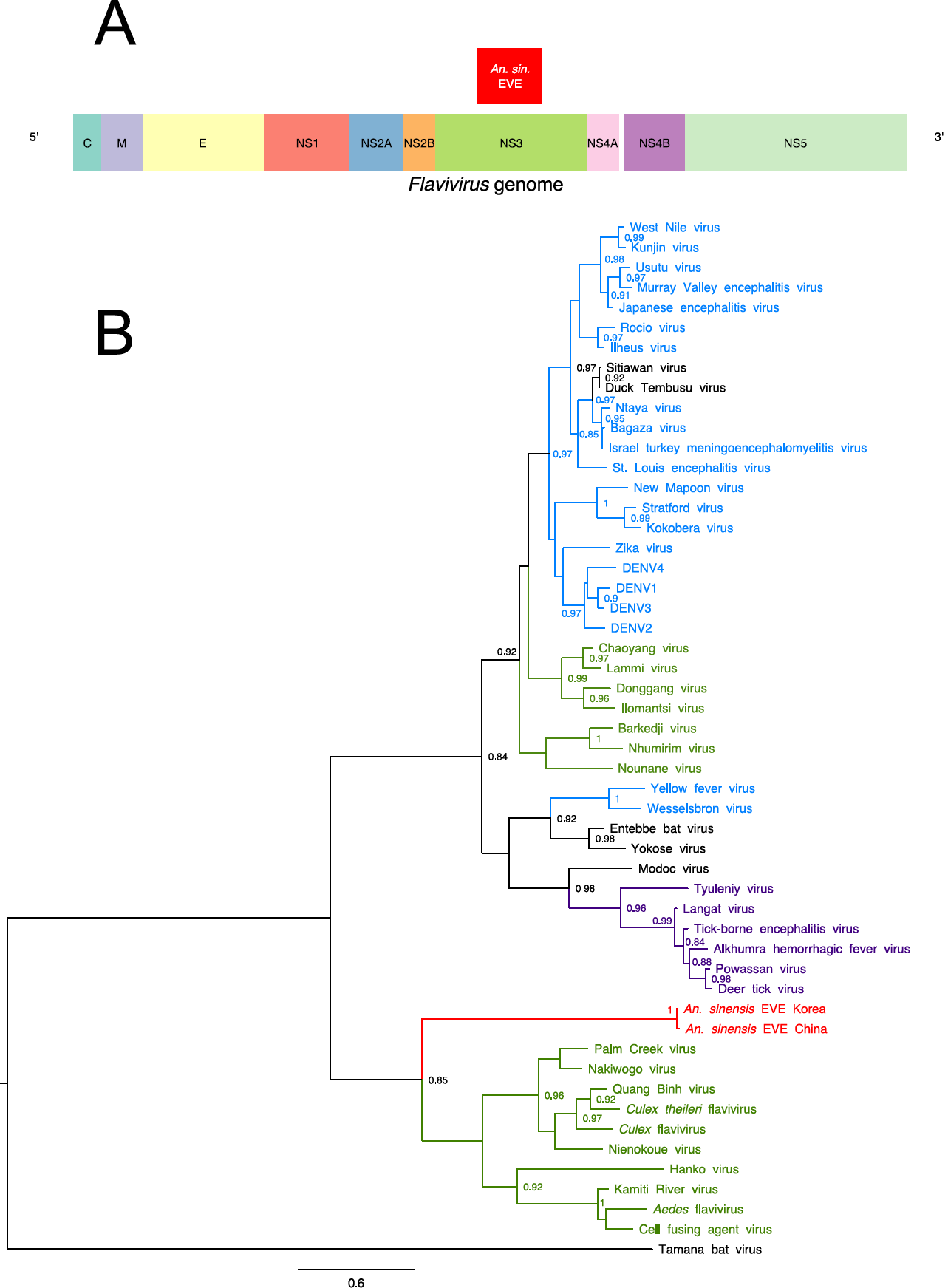
Discovery of a flavivirus-derived EVE in *An. sinensis*. (A) EVE location in a generic *Flavivirus* genome. Positioning is based on the genome sequence of *Culex* flavivirus (GenBank accession no. JQ308188). C=capsid protein, E=envelope glycoprotein, M=membrane glycoprotein, NS1=non-structural glycoprotein 1; NS2A=non-structural protein 2A; NS2B= non-structural protein 2B; NS3=non-structural protein 3 (protease/helicase); NS4A=non-structural protein 4A; NS4B=non-structural protein 4B; NS5=non-structural protein 5 (RNA-dependent-RNA polymerase). (B) Phylogenetic relationships of the newly discovered *An. sinensis* flavivirus-derived EVEs with exogenous flaviviruses. Maximum likelihood trees were constructed based on the translated EVE sequence. Clades are color-coded according to known host specificity: green, ISFs; purple, tick-borne arboviruses; black, ‘not known vector’ (vertebrate specific); blue, mosquito-borne arboviruses; red: EVEs. Scale bar indicates the number of substitutions. Node values represent Shimodaira-Hasegawa (SH)-like branch support (only values > 0.8 are shown).

The *An. sinensis* supercontig containing the EVE is 2,797 bp long for the Korean strain and 7,380 bp long for the Chinese strain. Analyses of flanking regions revealed the presence of another EVE in the same orientation, upstream of the flavivirus-derived EVE in both the Korean and the Chinese strains. The closest BLAST hit of this additional EVE is Xincheng mosquito virus (43% and 42% amino-acid identity for the Korean and Chinese strains, respectively), an unclassified, negative-sense, single-stranded RNA virus detected in a pool of Chinese mosquitoes including *An. sinensis* specimens (Table S3). BLAST and conserved domain search identified a class II Mariner-like transposase close to a mariner mos1 element, approximately 1,000 bp downstream of the flavivirus-derived EVE (Table S3). This was only the case for the Chinese strain because the supercontig of the Korean strain was not long enough.

Presence of the *An. sinensis* EVE was verified *in vivo* by PCR on genomic DNA from the Korean strain (Figure 4A), followed by amplicon sequencing to confirm identity. The *An*. *sinensis* EVE was found in both male and female genomic DNA, and was transcriptionally expressed, although less abundantly than the *An. sinensis* EVE, for all combinations of sex and development stages tested, especially in L_4_ larvae (Figure 4B). Low expression observed for the *An. sinensis* EVE is consistent with barely detectable transcriptional activity in published RNA-seq data (Figure S1B).

**Figure 4:**
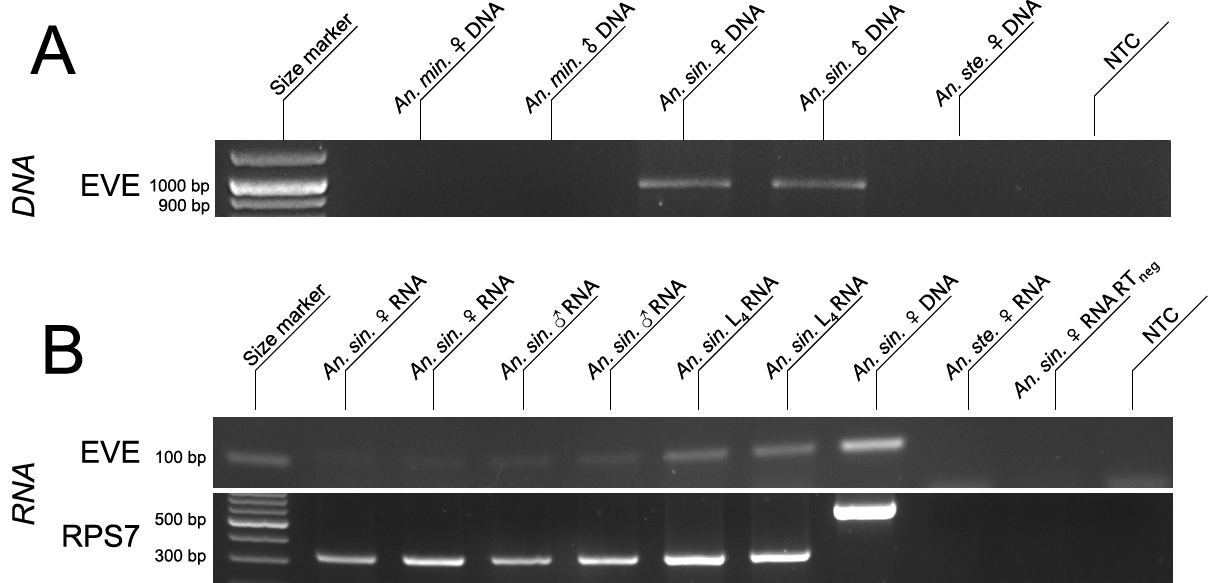
*In vivo* detection of the *An. sinensis* flavivirus-derived EVE. (A) EVE detection in genomic DNA from the Korean strain of *An. sinensis*. Lane 1: size marker; lane 2: amplified genomic DNA from a pool of 10 *An. minimus* adult females; lane 3: amplified genomic DNA from a pool of 10 *An. minimus* adult males; lane 4: amplified genomic DNA from a pool of 10 *An. sinensis* adult females; lane 5: amplified genomic DNA from a pool of 10 *An. sinensis* adult males; lane 6: amplified DNA from a pool of 10 *An. stephensi* females; lane 7: no template control (NTC). (B) EVE detection in total RNA from the Korean strain of *An. sinensis*. Lane 1: size marker; lanes 2 and 3: amplified cDNA from pools of 5 adult females; lanes 4 and 5: amplified cDNA from pools of 5 adult males; lanes 6 and 7: amplified cDNA from pools of 5 L_4_ larvae; lane 8: amplified DNA from a pool of 10 females; lane 9: amplified cDNA from a pool of 5 *An. stephensi* females; lane 10: DNA contamination control (no reverse transcription) using the same pool of 5 adult females as lane 2; lane 11: no template control (NTC). First row: EVE; second row: RPS7 (control gene). The RPS7 target DNA sequence includes an intron, so that DNA contamination is expected to result in a larger PCR product.

## Discussion

ISFs have attracted substantial interest in recent years after some of them were shown to enhance or suppress the replication of medically important flaviviruses in co-infected mosquito cells (Blitvich and Firth 2015). Over a dozen of ISFs have been formally identified to date, mainly in *Aedes* and *Culex* genera of the Culicinae subfamily (Blitvich and Firth 2015). ISFs were also reported in *Anopheles* mosquitoes of the Anophelinae subfamily (Aranda et al. 2009; Zuo et al. 2014; Liang et al. 2015). However, these *Anopheles*-associated ISFs are thought to infect a broad range of hosts including several mosquito species, mainly in the *Culex* genus, and are phylogenetically related to *Culex*-associated ISFs. Therefore, it is unclear whether *Anopheles* mosquitoes are true natural hosts of flaviviruses. Detection of ISFs in field-caught mosquitoes could result from incidental infection, or from a laboratory artifact. In this study, we discovered flavivirus-derived EVEs in the genomes of two *Anopheles* species. Phylogenetic analyses indicated that both EVEs are related to ISFs but belong to a clade that is distinct from *Aedes*-associated and *Culex*-associated ISFs.

Presence of flavivirus-derived EVEs in *Anopheles* genomes supports the hypothesis that *Anopheles* mosquitoes are natural hosts of flaviviruses. Endogenization of non-retroviral RNA viruses is unlikely to occur in the absence of recurrent host-virus interactions over a long evolutionary time scale. Endogenization requires reverse transcription, germ line integration and fixation in the host population, three steps whose combined frequency is exceedingly rare (Holmes 2011; Aiewsakun and Katzourakis 2015). The species-wide frequency of the *An*. *minimus* EVE is unknown because our *in silico* and *in vivo* analyses were based on the same mosquito strain. Presence of the *An. sinensis* EVE in two mosquito strains from different geographical locations, however, suggests that it could be fixed at the species level. Thus, our discovery of flavivirus-derived EVEs in *Anopheles* genomes is consistent with a long-lasting host-virus interaction between flaviviruses and mosquitoes of the Anophelinae subfamily.

ISFs are thought to be mainly maintained through vertical transmission from an infected female to its offspring (Blitvich and Firth 2015). Vertical transmission is likely to favor co-divergence of pathogens and hosts (Jackson and Charleston 2004), as illustrated by the existence of *Aedes*-associated and *Culex*-associated clades of ISFs (Moureau et al. 2015). Although extrapolation is limited by the scarcity of data on ISF host range and diversity, phylogenetic position of *Anopheles*-associated ISFs as sister to all other known ISFs is consistent with the co-divergence hypothesis. During the evolutionary history of mosquitoes, the Anophelinae diverged from the Culicinae prior to the separation of *Culex* and *Aedes* genera (Reidenbach et al. 2009). Further investigations will be necessary to determine whether an *Anopheles*-associated clade of exogenous ISFs exists, or existed.

Non-retroviral EVEs are thought to be generated by interaction of exogenous viruses with endogenous retro-elements, either with or without long terminal repeats (LTR) (Holmes 2011). The short size of the supercontigs containing the *An. minimus* and *An. sinensis* EVEs limited our ability to investigate the integration mechanism(s). Another EVE sequence that we identified close to the flavivirus-derived EVE in *An. sinensis* may point to an EVE hotspot.

Sequence conservation of the *An. minimus* EVE (i.e., absence of stop codon across 1,881 bp) is consistent with a recent integration or a more ancient integration followed by non-neutral evolution. Our observation that the *An. minimus* EVE is abundantly transcribed may reflect a selective advantage for the host (Holmes 2011). Transcriptionally active EVEs have been suggested to confer protection or tolerance against related exogenous viruses (Flegel 2009; Holmes 2011; Aswad and Katzourakis 2012; Bell-Sakyi and Attoui 2013; Fujino et al. 2014). Despite the lack of empirical evidence so far, flavivirus-derived EVEs could contribute to antiviral immunity and arbovirus vector competence in mosquitoes.

## Acknowledgements

This work was supported by the French Government’s Investissement d’Avenir program Laboratoire d’Excellence Integrative Biology of Emerging Infectious Diseases grant ANR-10-LABX-62-IBEID, and the City of Paris Emergence(s) program in Biomedical Research. S.L. was supported by a doctoral fellowship from University Pierre and Marie Curie. The funders had no role in study design, data collection and interpretation, or the decision to submit the work for publication.

The authors thank Inge Holm and Guillaume Carissimo for providing *Anopheles stephensi* nucleic acids, Albin Fontaine for help with the *in silico* screen, Davy Jiolle and Elliott Miot for technical assistance, Clément Gilbert for advice and critical reading of an earlier version of the manuscript, and members of the Lambrechts lab for insightful comments and discussions.

## Supporting information

**Table S1:** Names and accession numbers of flavivirus polyprotein sequences used as queries for the *Anopheles* genome screen and included in the EVE phylogenetic analyses.

**Table S2:** PCR primers and reaction conditions used to amplify EVE and RPS7 targets from DNA or cDNA templates.

**Table S3:** Other genetic integrations detected in the *An. sinensis* genome supercontigs containing the flavivirus-derived EVE.

**Figure S1:** Sequencing coverage of (A) *An. minimus* and (B) *An. sinensis* EVEs in published RNA-seq experiments. For each experiment, the Sequence Read Archive (NCBI) accession number is indicated above the graph.

